# The representation of regularity and randomness in auditory cortex of awake rats

**DOI:** 10.1101/2025.04.09.647967

**Authors:** Sapir Bashari, Maciej M. Jankowski, Israel Nelken

## Abstract

Auditory cortex possesses a remarkable ability to discriminate between tone sequences with different levels of statistical regularities. To examine this phenomenon at the level of single neurons, we recorded responses from auditory cortex of awake rats. The rats passively listened to multitone sequences composed of 4, 5 or 6 tones. The tones could be presented in periodic sequences with a fixed, repeated cycle; in a fully random condition; or in an intermediate condition in which cycles were maintained but tones were randomly permuted within each cycle (random cycle condition). Units showed a continuum of preferences between order-preferring neurons and randomness-preferring neurons. Unexpectedly, we found sensitivity to the position of sounds in the cycle (‘phase modulation’). The strongest and most consistent phase modulations were in the responses to the random cycle condition. Finally, we show how such sensitivities may emerge from a simple, biologically-plausible computation.

## Introduction

Responses to sounds in auditory cortex depend not only on the physical characteristics of sounds but also on their spectral and temporal context. In particular, auditory cortex is sensitive to statistical regularities in incoming sound sequences. Sensitivity to statistical regularities is the hallmark of stimulus-specific adaptation (SSA), the decrease in neural responses to a common stimulus that generalizes only partially to other, rare stimuli ^1^: neurons in auditory cortex exhibit a stronger response to a sound when it occurs with a low probability compared to the same sound occurring with a higher probability^2–4^. The larger responses to rare and deviant sounds have been shown in many species^1,5–11^, in awake^12,13^ as well as in anesthetized animals. SSA is believed to be the neural correlate of the deviance sensitivity of the middle latency responses (MLRs) recorded in human subjects^14,15^. However, the sensitivity to temporal regularities extends beyond SSA: for example, neurons in the auditory cortex of anesthetized rats responded more strongly to the same tones when they appeared in random sequences compared to regular sequences, even when their probability of occurrence was the same^16^.

Remarkably, studies in humans that used multitone sequences^17,18^ found that the strength of the response in the auditory cortex to such sequences is also related to the predictability of the tones in the sequence. In contrast in animals, in these experiments, more predictable sequences produced larger brain signals.

In this study we used multitone sequences spanning the range from fully random to periodic, recording neuronal responses in auditory cortex of awake rats. We show an exquisite sensitivity to the level of regularity of the sequence, suggest how common phenomena may underlie the opposite results in humans and animal models, and show how such sensitivity can emerge from a simple, biologically plausible computation.

## Results

### Multitone sequences

Every sequence had 60 tones (50 ms long, 10 ms rise fall times), presented without gaps between them. The sequences were generated by selecting T (T=4, 5, or 6) frequencies from 20 possible values (4-40 kHz, 6 frequencies/octave) and the same frequencies were used throughout the sequence. For every selection of T tones, we generated sequences in three conditions – fixed cycle, random cycle, and full random - differing in the degree of order in their periodic structure (cycles). Fixed cycle sequences were generated by selecting a random permutation of the T selected frequencies and then using that same permutation in every cycle of the sequence (Figure 1A). In the fixed cycle sequences, cycles had a clear meaning: the same tone frequencies were repeated in the same order in each cycle. Random cycle sequences were generated by selecting a new random permutation of the T frequencies for each cycle. In this condition, the cycles were defined by the rule that each of the T tones appeared only once within each cycle. Full random sequences consisted of T selected frequencies that appeared the same number of times in the sequence (equal to the number of cycles in the fixed cycle and random cycle conditions), with no other constraint on the order in which they occurred in the sequence.

**Figure 1.**
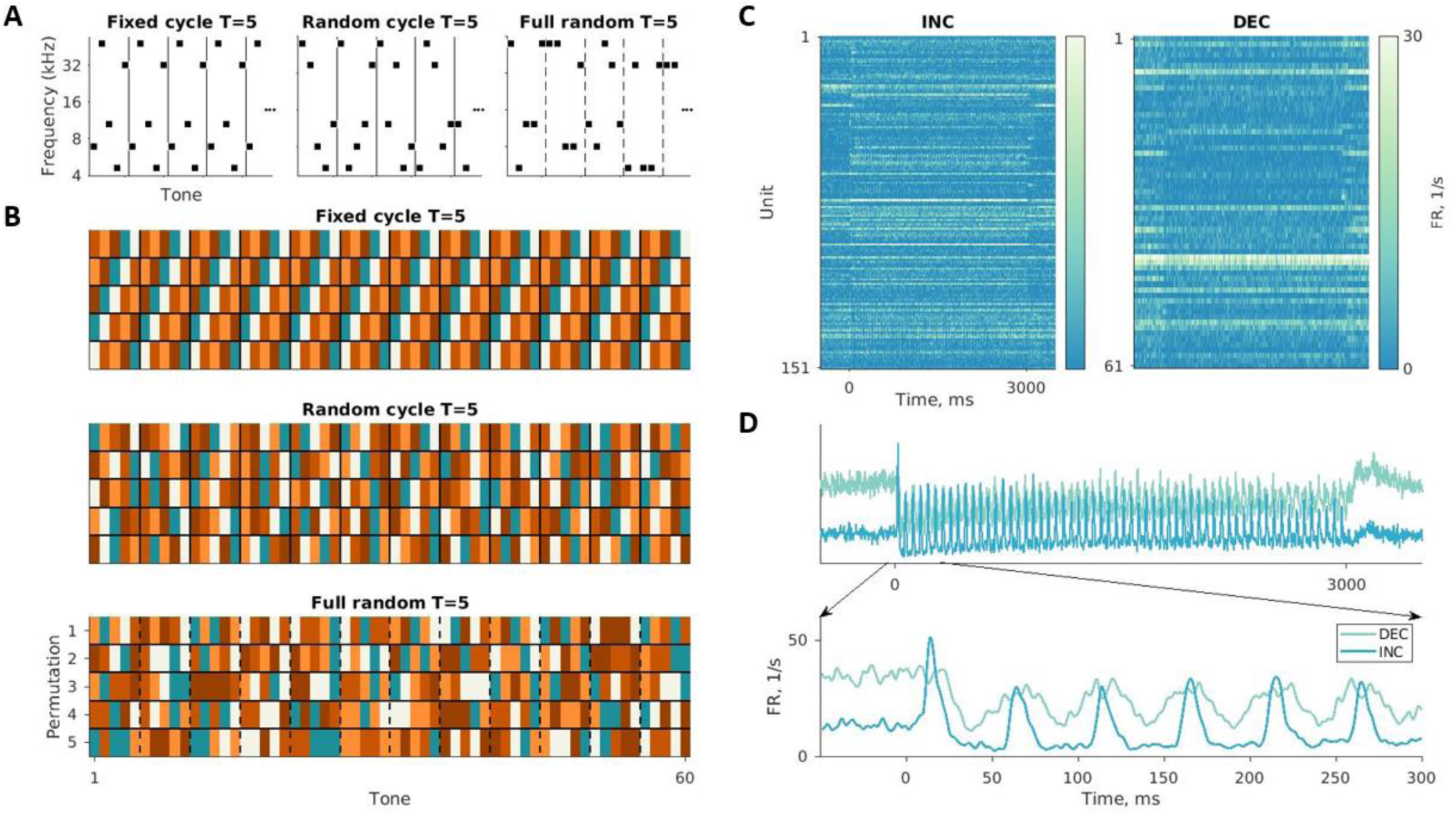
Multitone sequences A. Examples of multitone sequences with T=5. The X-axis represents the position of the tone within the sequence, and the Y-axis is (log) frequency. The lines separate between cycles. In the full random case, cycles are nominal only. B. Example of all the sequences generated for 5 selected tones. For every choice of T tones there were T sequences in each condition. The colors represent the frequencies, the x-axis is the index of tones in the sequence and the y-axis represents the T sequences generated for each of the frequency choices. For fixed cycle sequences, cyclic permutations were used. For random cycle sequences, the permutations were random within each cycle in all sequences. For full random sequences, random permutations of the entire sequences were used. Black lines separate between the cycles (dashed lines are used for the full random cycle sequences, to emphasize the lack of real cycle structure in this case). C. The responses of all "INC" and "DEC" units in one recording session. Each row represents the response of one unit. The color represents the firing rate of that unit, averaged over all sequences. D. Mean responses over all sequences of an exemplary "INC" unit and an exemplary "DEC" unit recorded in the same session. "INC" units showed an increase, and “DEC” units a decrease, in their average firing rate relative to baseline in response to the first tone of the sequence. Here and elsewhere, firing rate traces were smoothed with a Hamming window of length 5 ms.

The sequences had 15, 12 or 10 cycles when T was 4, 5 and 6 tones/cycle respectively. Each tone can be identified by its location in the sequence (1-60), but also by the cycle in which it occurs (1-15 for T=4, 1-12 for T=5, and 1-10 for T=6) as well as its position within the cycle, which we refer to as the ‘phase’ of the tone in the cycle (1,2,…,T).

Multiple sequences were generated with the same tone frequencies in order to control for the effects of frequency sensitivity on the structure of the neuronal responses (Figure 1B, see Methods for details).

The data were collected from four awake non-behaving rats using Neuropixels electrodes. The data shown in the results section consists of the responses to sets of T=4/5/6 sequences collected from two rats. From each of these two rats we also recorded the responses to another set of T=5 sequences with only two or three selections of 5 frequencies, but many more permutations of those; thus, the use of individual frequencies was more balanced than the main data sets. In addition, responses to another set of T=5 sequences in which the frequencies were less balanced were collected in two more rats (see Methods). The analysis of these three additional data sets is presented in the supplemental figures. The results for the T=5 sequences were reproduced in the additional datasets.

The responses were quantified by the number of spikes emitted in 50 ms time windows, corresponding to the durations of the individual tones in the sequences (except when stated otherwise). We analyzed units that had high enough, stable firing rate and also showed a significant difference between baseline activity and the response to the first tone in the sequences (inhomogeneous Poisson test with p<0.05). In the main data, 4404 units (out of 41007 recorded units) fulfilled these criteria (see Methods for details). We distinguished between units that showed a significant increase in their firing rate to the first tone compared to baseline (“INC” units) and those that showed a decrease ("DEC" units; Figures 1C and 1D).

The responses to the first T tones in each sequence were not included in the analysis because during the first cycle there is still no difference between fixed cycle and random cycle sequences. As opposed to the first cycle of these conditions, the first T tones in full random sequences could in principle provide hints for the type of the sequence (about 95% of the full random sequences contained tone repetitions in their first T tones); however, to be consistent with the other two conditions, we ignored the first T tones in the full random condition as well.

### A continuum of order- and randomness-preferring units

We decomposed the responses to each tone into a sum of contributions of tone frequency and condition using linear mixed-effects (LME) models (see Methods). Averaged over all units and tones, the population responses depended significantly on the condition in all sets except for Set3 (Table 1).

**Table 1.**
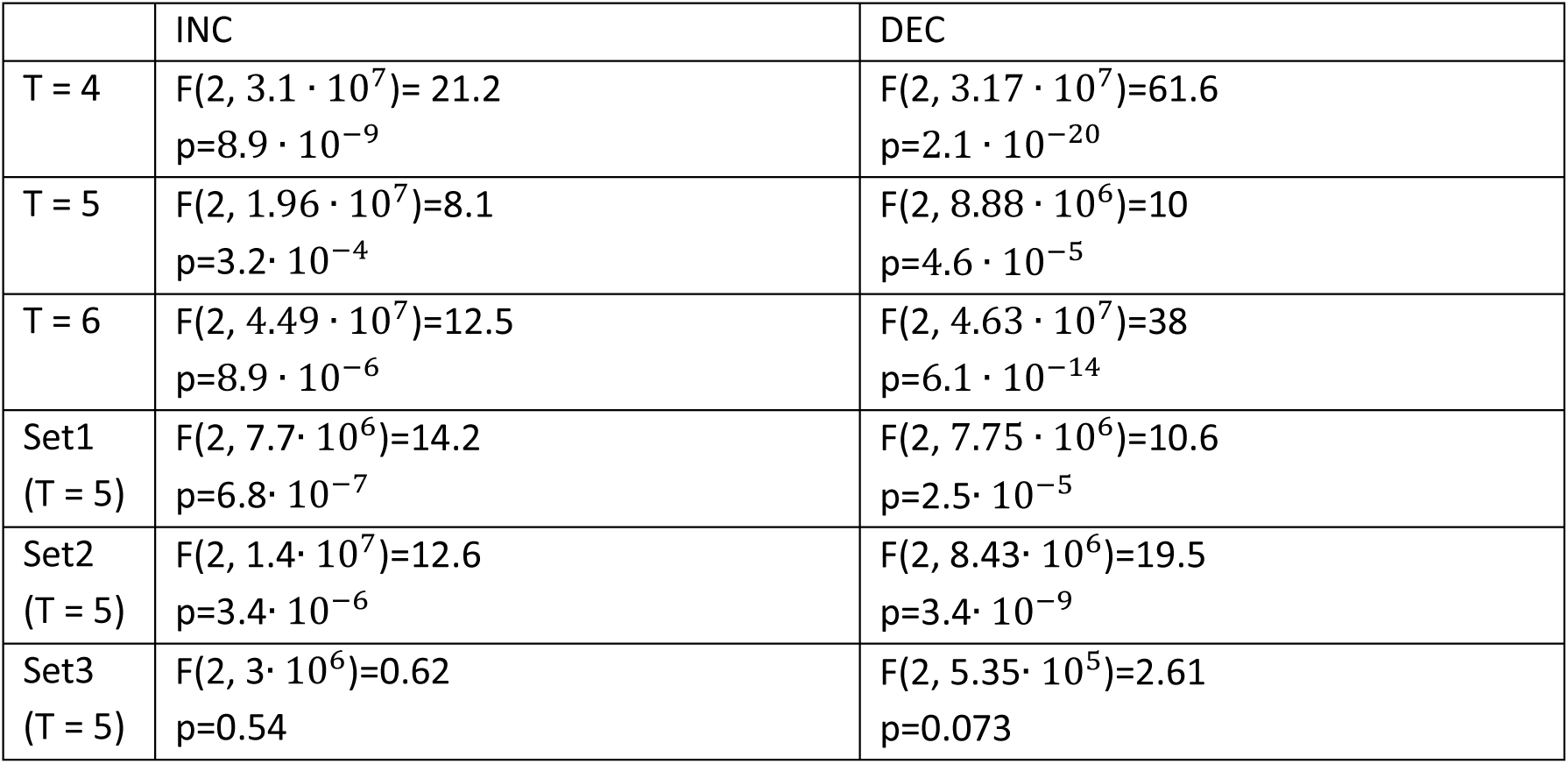
Dependence on condition Tests for the fixed effect of condition for all data sets and unit types.

|  | INC | DEC |
| --- | --- | --- |
| T = 4 | $F(2, 3.1 \cdot 10^7) = 21.2$<br>$p = 8.9 \cdot 10^{-9}$ | $F(2, 3.17 \cdot 10^7) = 61.6$<br>$p = 2.1 \cdot 10^{-20}$ |
| T = 5 | $F(2, 1.96 \cdot 10^7) = 8.1$<br>$p = 3.2 \cdot 10^{-4}$ | $F(2, 8.88 \cdot 10^6) = 10$<br>$p = 4.6 \cdot 10^{-5}$ |
| T = 6 | $F(2, 4.49 \cdot 10^7) = 12.5$<br>$p = 8.9 \cdot 10^{-6}$ | $F(2, 4.63 \cdot 10^7) = 38$<br>$p = 6.1 \cdot 10^{-14}$ |
| Set1<br>(T = 5) | $F(2, 7.7 \cdot 10^6) = 14.2$<br>$p = 6.8 \cdot 10^{-7}$ | $F(2, 7.75 \cdot 10^6) = 10.6$<br>$p = 2.5 \cdot 10^{-5}$ |
| Set2<br>(T = 5) | $F(2, 1.4 \cdot 10^7) = 12.6$<br>$p = 3.4 \cdot 10^{-6}$ | $F(2, 8.43 \cdot 10^6) = 19.5$<br>$p = 3.4 \cdot 10^{-9}$ |
| Set3<br>(T = 5) | $F(2, 3 \cdot 10^6) = 0.62$<br>$p = 0.54$ | $F(2, 5.35 \cdot 10^5) = 2.61$<br>$p = 0.073$ |

Figure 2A shows the average response of INC and DEC units for different values of T in the main data. To emphasize the effect of condition, Figure 2B shows the difference between the response in each condition and their average. The peak responses in the full random condition were on average the largest, for both INC and DEC units, and for all cycle lengths.

**Figure 2.**
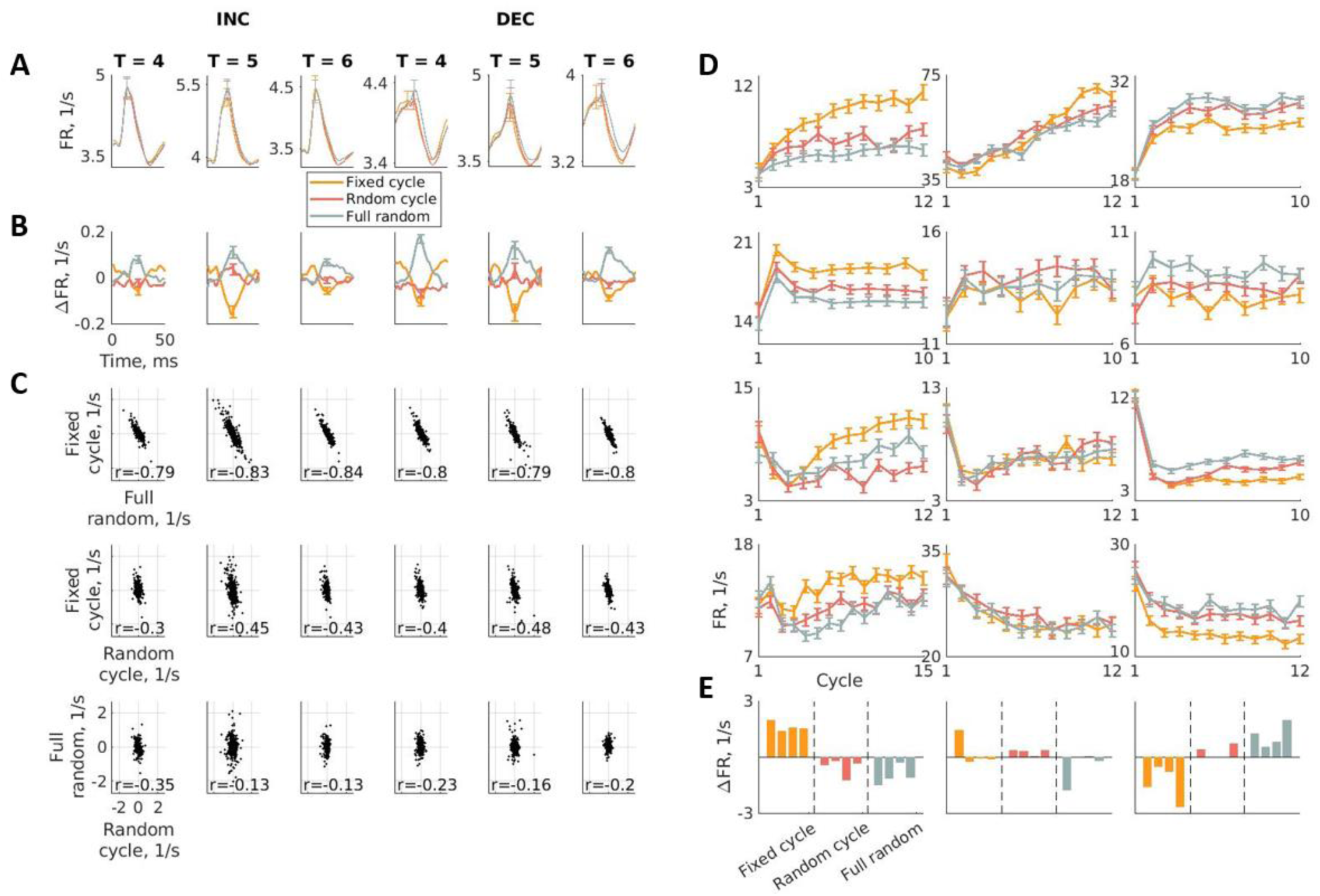
Dependence on condition and cycle A. The average responses of all auditory units to the tones in the three conditions. The three left panels show the average responses of INC units in all conditions and the three right panels of DEC units. B. The difference between the mean tone response in each condition and their overall mean. C. Scatter plots of the unit-specific contributions in the three conditions. Note the differences between the sizes of the main effects of condition presented in B (order of 0.1/s) and that of the unit-specific contributions shown in C (order of 1/s, an order of magnitude larger). See also Figure S1. The correlation coefficients are presented at the bottom of each panel. See also Figure S2. D. Examples of order-preferring units (left column), randomness preferring units (right column) and less selective units (middle column). The responses of each unit were averaged over all phases within each cycle, and over all sequences of the corresponding condition. They are plotted as a function of the cycle number (here, including the first cycle). See also Figure S3 and Figure S4. E. Examples of unit-specific contributions for the three conditions. Each panel shows the contributions of the units in the corresponding column in D. The dashed lines separate the three conditions. Fixed cycle in orange, random cycle in red and full random in gray. Within each condition, the order of units (left to right) corresponds to their order in D (top to bottom).

Similarly, the peak responses were on average smallest in the fixed cycle condition. The responses in the random cycle condition were intermediate.

Importantly, responses of individual units were often very different from the population averages. Indeed, the unit-specific contributions for condition significantly improved the fit of the models to the data (Table 2). In all sets, the standard deviations of the unit-specific contributions for condition (measuring the variability of the effects across units) were much larger than the main effects (that roughly represent the mean over all units; Figure S1). In consequence, the condition-dependence of individual units could be very different from the population average. Remarkably, the unit-specific contributions showed negative correlations between conditions (Figure 2C for the main data; Figure S2 for the additional datasets). The strongest negative correlations were found between the unit-specific contributions for the fixed cycle and full random conditions - the most regular and most random conditions - with weaker correlations of either of them with the unit-specific contributions of the random cycle condition. Indeed, for about 50% of the units, the unit-specific contributions for the random cycle condition were intermediate between those of the fixed cycle and full random conditions, accounting for this correlation structure (the expected fraction was 33%; Table 3; Fisher’s exact test, p<=0.0001 in 10/12 cases).

**Table 2.** The unit-specific contributions for condition were highly significant Likelihood ratio tests between LME models with and without unit-specific random contributions for condition. All p=0.

|  | T = 4 | T = 5 | T = 6 | Set1 (T=5) | Set2 (T=5) | Set3 (T=5) |
| --- | --- | --- | --- | --- | --- | --- |
| INC | $\chi^2(3)= 8914.4$ | $\chi^2(3)= 12364$ | $\chi^2(3)= 12305$ | $\chi^2(3)= 2949.6$ | $\chi^2(3)= 9027.2$ | $\chi^2(3)= 1891.1$ |
| DEC | $\chi^2(3)= 9843.5$ | $\chi^2(3)= 5510.3$ | $\chi^2(3)= 9578.9$ | $\chi^2(3)= 4030.8$ | $\chi^2(3)= 5233.6$ | $\chi^2(3)= 231.92$ |

**Table 3.** The units-specific contributions to the random cycle condition tended to be intermediate between those for the fixed cycle and full random conditions The cases in which the unit-specific factors for the random cycle condition were intermediate between those of the fixed cycle and full random conditions (expected fraction: 33%), as well as their subdivisions to the two possible cases (fixed cycle maximal or full random maximal; expected fraction: 16%). Fisher’s exact test.

| Set |  | Number of cases | Random cycle were intermediate |  | % fixed cycle < random cycle < full random | % full random < random cycle < fixed cycle |
| --- | --- | --- | --- | --- | --- | --- |
|  |  |  | % of cases | P value |  |  |
| T=4 | INC | 539/1200 | 0.45 | P<0.0001 | 0.22 | 0.23 |
|  | DEC | 589/1226 | 0.48 | P<0.0001 | 0.25 | 0.23 |
| T=5 | INC | 670/1350 | 0.5 | P<0.0001 | 0.23 | 0.27 |
|  | DEC | 303/628 | 0.48 | P<0.0001 | 0.2 | 0.28 |
| T=6 | INC | 587/1200 | 0.49 | P<0.0001 | 0.24 | 0.25 |
|  | DEC | 627/1226 | 0.51 | P<0.0001 | 0.21 | 0.3 |
| Set1<br>(T=5) | INC | 177/479 | 0.37 | P=0.14 | 0.16 | 0.2 |
|  | DEC | 233/483 | 0.48 | P<0.0001 | 0.27 | 0.21 |
| Set2<br>(T=5) | INC | 297/587 | 0.51 | P<0.0001 | 0.22 | 0.29 |
|  | DEC | 174/352 | 0.49 | P<0.0001 | 0.2 | 0.3 |
| Set3<br>(T=5) | INC | 230/382 | 0.6 | P<0.0001 | 0.28 | 0.32 |
|  | DEC | 31/69 | 0.45 | p=0.11 | 0.26 | 0.19 |

Figure 2D shows the responses of exemplary units with different preferences to order and randomness. Order-preferring units are presented in the left column, randomness-preferring units in the right column, and less selective units in the middle column. The unit-specific contributions for the three conditions are shown in Figure 2E. In the left column, the unit-specific contributions for the fixed cycle condition (orange) are positive, for the full random condition (grey) negative, and for the random cycle condition (red) they are intermediate; in the right column, the preference is opposite; while in the central column, these contributions are small in all conditions.

Thus, in contrast with the anesthetized auditory cortex, were the large majority of neurons respond more strongly to tones in random than in regular sequences^16^, units in the awake auditory cortex showed a continuum of preferences between order-preferring neurons (whose responses to the fixed cycle condition were stronger, and to the full random condition weaker), and randomness-preferring neurons (whose responses to the fixed cycle condition were weaker, and to the full random condition stronger).

We analyzed the modulation pattern of the firing rate during the 50 ms of a tone presentation. Within this period, the average responses started at baseline, increased to their maximum, and decrease back to baseline. Interestingly, the dependence on condition differed at baseline and at the peak. We therefore analyzed separately the peak response (the spike count between 11 to 40 ms after each tone onset) and the baseline firing rates (1- 10 and 41-50 ms after sound onset, at the beginning and the end of each 50 ms tone presentation).

As expected, the peak responses in the full random condition had the largest averages, in the fixed cycle condition the smallest, and in the random cycle condition intermediate averages (Table 4). In contrast with the peak responses, during the baseline window, firing rates were on average largest in the fixed cycle condition in most cases (Table 4). Thus, while peak rates were on average smaller in the fixed cycle condition than in the full random and random cycle conditions, they occurred over a baseline that tended to be larger on average in the fixed cycle condition.

**Table 4.** Condition modulation patterns of the firing rate during a tone presentation Significance of the differences between the average population responses to tones played in the three conditions in the peak response window (11-40 ms after tone onset) and the baseline window (1-10 and 41-50 ms after tone onset), derived using permutation tests. All p-values are two-sided; in all significant cases, the sign of the difference was the same and is specified at the top of each column.

| Set |  | Peak |  |  | Baseline |  |  |
| --- | --- | --- | --- | --- | --- | --- | --- |
|  |  | Fixed cycle < full random | Fixed cycle < random cycle | random cycle < full random | Fixed cycle > full random | Fixed cycle > random cycle | Random cycle < full random |
| T=4 | INC | P=0.001 | n.s | P=0.001 | P=0.001 | P=0.001 | P<0.014 |
|  | DEC | P=0.001 | n.s | P=0.001 | P=0.001 | P=0.001 | n.s |
| T=5 | INC | P=0.001 | P=0.001 | P<0.003 | P<0.008 | P<0.003 | n.s |
|  | DEC | P=0.001 | P<0.003 | P=0.001 | n.s | P<0.009 | n.s |
| T=6 | INC | P=0.001 | n.s | P=0.001 | n.s | P<0.012 | n.s |
|  | DEC | P=0.001 | P<0.002 | P=0.001 | n.s | n.s | P<0.01 |
| Set1 (T=5) | INC | P=0.001 | P=0.001 | P=0.001 | P<0.015 | n.s | P<0.013 |
|  | DEC | n.s | n.s | n.s | P=0.001 | P=0.001 | n.s |
| Set2 (T=5) | INC | P=0.001 | P=0.001 | P=0.001 | n.s | P<0.003 | P=0.001 |
|  | DEC | P=0.001 | n.s | P=0.001 | n.s | P<0.021 | P=0.001 |
| Set3 (T=5) | INC | P<0.014 | P<0.01 | n.s | P=0.001 | P=0.001 | n.s |
|  | DEC | n.s | n.s | P<0.011 | n.s | n.s | n.s |

### A large variability of effects of cycle

To study the dependence of neural responses on cycle we fitted LME models for each condition separately. We decomposed the responses to each tone into a sum of contributions of frequency, cycle, and phase modulation using LME models (see Methods).

In most of the sets the average population responses did not show significant main effect of cycle, so the average population response remained largely stable along the sequences.

However, the unit-specific contributions significantly improved the fit of the models (Table 5). Indeed, the standard deviations of the unit-specific contributions were much larger than the main effects (Figure S3).

**Table 5.** The unit-specific contributions for cycle were highly significant Likelihood ratio tests between LME models with and without unit-specific random factors for cycle. All p=0.

|  |  | fixed cycle | random cycle | full random |
| --- | --- | --- | --- | --- |
| T=4 | INC | $\chi^2(2) = 2924.5$ | $\chi^2(2) = 2891.6$ | $\chi^2(2) = 2437.3$ |
| | DEC | $\chi^2(2) = 3099.1$ | $\chi^2(2) = 1694$ | $\chi^2(2) = 837.81$ |
| T=5 | INC | $\chi^2(2) = 14256$ | $\chi^2(2) = 6354.2$ | $\chi^2(2) = 5001.5$ |
| | DEC | $\chi^2(2) = 4466.9$ | $\chi^2(2) = 2164.6$ | $\chi^2(2) = 2068.7$ |
| T=6 | INC | $\chi^2(2) = 2215.3$ | $\chi^2(2) = 1636.5$ | $\chi^2(2) = 1936.9$ |
| | DEC | $\chi^2(2) = 1684.6$ | $\chi^2(2) = 1028.5$ | $\chi^2(2) = 978.7$ |
| Set1<br>(T=5) | INC | $\chi^2(2) = 610.27$ | $\chi^2(2) = 309.1$ | $\chi^2(2) = 280.7$ |
| | DEC | $\chi^2(2) = 520.28$ | $\chi^2(2) = 172.18$ | $\chi^2(2) = 94.61$ |
| Set2<br>(T=5) | INC | $\chi^2(2) = 3638.2$ | $\chi^2(2) = 1493.9$ | $\chi^2(2) = 1236.4$ |
| | DEC | $\chi^2(2) = 2542.5$ | $\chi^2(2) = 848.28$ | $\chi^2(2) = 776.12$ |
| Set3<br>(T=5) | INC | $\chi^2(2) = 454.24$ | $\chi^2(2) = 223$ | $\chi^2(2) = 425.77$ |
| | DEC | $\chi^2(2) = 192.46$ | $\chi^2(2) = 90.53$ | $\chi^2(2) = 79.85$ |

While responses sometimes decreased over cycles (Figure 2D, lower row), as expected in anesthetized animals, many units showed increase as a function of cycle (Figure 2D, upper row), stable firing rates as a function of cycle, or even more complex patterns of change along the sequence (Figure 2D, middle rows). These behaviors tended to be similar for all conditions in each unit (Figure 2D), and were reflected in high correlations between the respective unit-specific contributions (Figure S4).

### Dependence on the phase in the cycle

Our main interest in this study was the dependence of the neural responses on the phase (position of the tone within cycle). The phase modulation is described using a "phase function". For a cycle of length T, the phase function consists of T values, one for each phase in the cycle. To estimate a phase function, the responses to each phase were averaged over cycles (except the first one) in all sequences, separately for each condition. The phase function consists of the deviations of the average response at each phase from the mean over all phases. Figure 3A shows the responses of one unit in the three conditions averaged over cycles (bottom), together with the derived phase functions (top). When the responses to all phases is the same, the phase function is flat (and close to 0). This was the expected result for the responses in the full random condition, since there was no real cycle structure in the sequence. Figure 3D shows further examples of cycle-averaged responses of different units with significant phase modulation in the fixed cycle condition (left column) and in the random cycle condition (right column), for sequences with T=4, 5 and 6.

**Figure 3.**
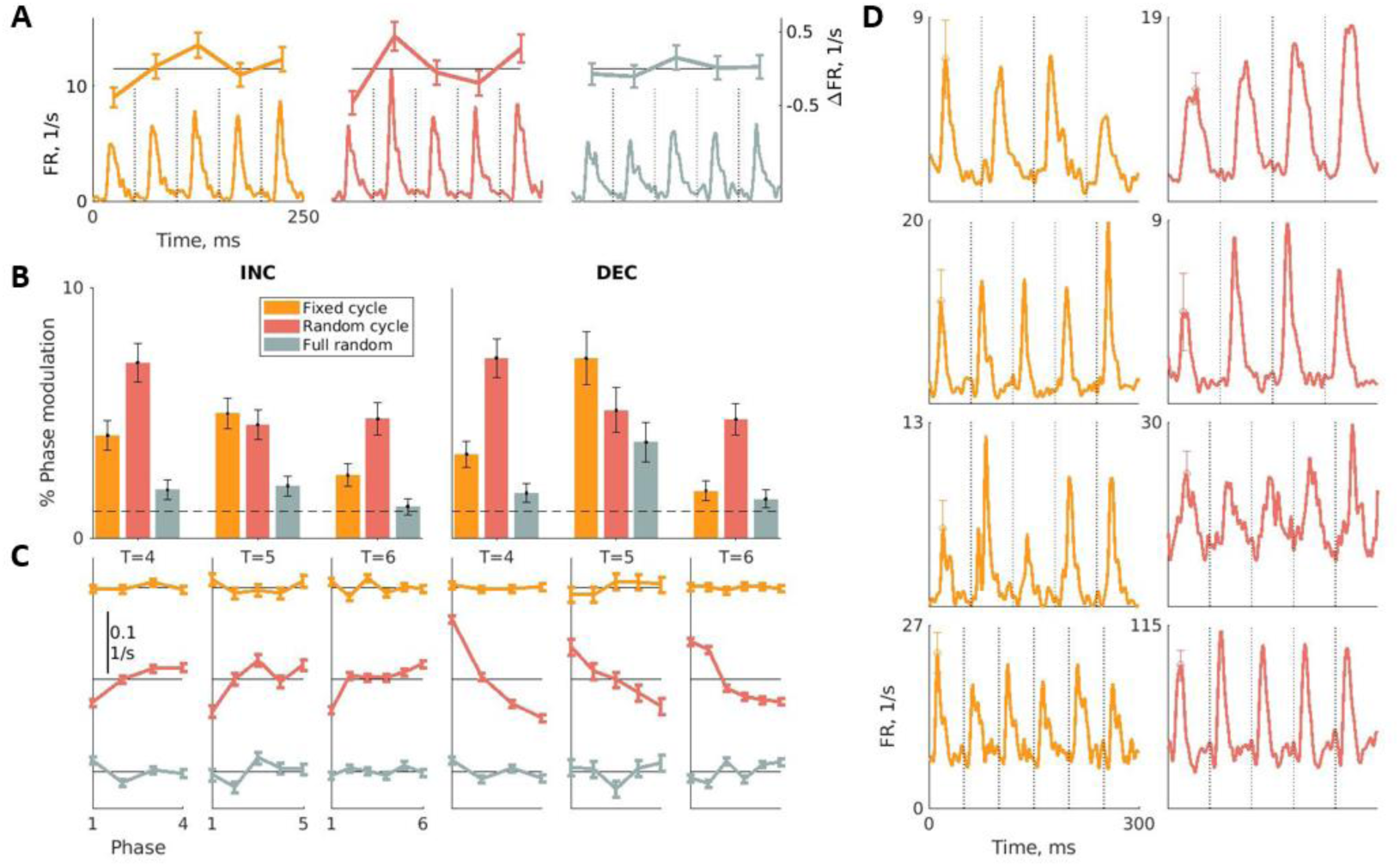
Dependence on the phase in the cycle A. Example of the responses of one unit averaged over all cycles of the T=5 sequences in three conditions (bottom) and the derived phase functions (top). The dashed lines mark the onset of successive tones and therefore appear every 50 ms. B. The percentage of units that showed significant phase modulation in each condition. The dashed line marks the expected value under the null hypothesis. See also Figure S5A and Figure S6. C. The mean population phase functions. See also Figure S5B and Figure S7. D. Cycle-average responses of units with significant phase modulation for fixed cycle sequences (left column) and for random cycle sequences (right column) and for T=4, T=5 and T=6. Each panel shows responses of a different unit.

For determining the significance of phase modulation, we modeled the responses of each unit, separately for each condition, as a sum of the effects of cycle, phase, and tone frequency. The phase function was tested against the null hypothesis of no phase modulation using a conservative permutation test (see Methods). All the examples in Figures 3A and 3D, except for the responses to the full random condition in Figure 3A, had significant phase modulation. Figure 3B shows the percentage of units in the main data whose phase functions had a significant modulation in the three conditions (fixed cycle, random cycle and full random) for T = 4, 5, and 6, separated by the unit type (INC and DEC). Figure S5A shows the same information for the other data sets. The percentage of units that showed significant phase modulation was significantly different between all conditions in the main data as well as for the additional datasets, except for DEC units in Set3 (Table 6).

**Table 6.** The fraction of units that showed significant phase modulation differed between conditions 𝜒^2^ tests of the number of significant units in the three conditions against the null of a uniform distribution across conditions.

|  | INC | DEC |
| --- | --- | --- |
| T=4 | $\chi^2(2)=37.67$, $p=6.61 \cdot 10^{-9}$ | $\chi^2(2)=47.83$, $p=4.11 \cdot 10^{-11}$ |
| T=5 | $\chi^2(2)=17.64$, $p=1.48 \cdot 10^{-4}$ | $\chi^2(2)=7.05$, $p=0.029$ |
| T=6 | $\chi^2(2)=27.42$, $p=1.11 \cdot 10^{-6}$ | $\chi^2(2)=28.39$, $p=6.84 \cdot 10^{-7}$ |
| Set1 (T=5) | $\chi^2(2)=8.18$, $p=0.017$ | $\chi^2(2)=16.27$, $p=2.93 \cdot 10^{-4}$ |
| Set2 (T=5) | $\chi^2(2)=31.01$, $p=1.85 \cdot 10^{-7}$ | $\chi^2(2)=13.42$, $p=0.0012$ |
| Set3 (T=5) | $\chi^2(2)=12.8$, $p=0.0017$ | $\chi^2(2)=1.85$, $p=0.4$ |

The fraction of units that had significant phase modulation in the full random condition serves as the estimate for the false detection rate of our statistical test. In all sets, the percentage of units that showed a significant phase modulation in the full random condition was indeed the smallest, and in many cases significantly so (Table 7). Phase modulation in the full random condition, even when significant, tended to be weaker than in the fixed cycle and random cycle conditions (Figure S6). Unexpectedly, in most of the sets the percentage of significant units was the largest in the random cycle condition (Figure 3B and Figure S5A).

**Table 7.** The fraction of units that had significant phase modulation is consistently smallest in the full random condition Fisher’s exact test comparing the number of units that showed significant phase effect in the full random condition vs. the number of significant cases in the fixed cycle condition and in the random cycle condition.

| Set |  | fixed cycle - full random | random cycle - full random |
| --- | --- | --- | --- |
| T=4 | INC | p=0.0013 | p<0.0001 |
|  | DEC | p=0.0535 | p<0.0001 |
| T=5 | INC | p<0.0001 | p=0.0002 |
|  | DEC | p=0.0064 | p=0.1693 |
| T=6 | INC | p=0.017 | p<0.0001 |
|  | DEC | p=0.4352 | p<0.0001 |
| Set1<br>(T=5) | INC | P=0.0073 | P=0.3221 |
|  | DEC | P<0.0001 | P=0.0169 |
| Set2<br>(T=5) | INC | P=0.0417 | P<0.0001 |
|  | DEC | P=0.0034 | P<0.0001 |
| Set3<br>(T=5) | INC | P=0.0417 | P<0.0001 |
|  | DEC | P=0.3097 | P=0.1829 |

Figures 3C and S5B displays the population-averaged phase functions and their standard errors. To compare conditions, we used here effect sizes: the standard deviation of the average phase function (‘signal’) divided by the mean standard error (‘noise’). The effect size was largest in the random cycle condition for all cycle lengths, T = 4, 5, and 6 (Figure S7), for the populations of INC as well as DEC units, in the main data as well as the additional data sets. Remarkably, the average phase functions in the random cycle condition were consistent across cycle lengths - in the INC population, the responses tended to be somewhat smaller than average at the beginning of the cycle and somewhat larger than the average at the end of the cycle, and vice versa in the DEC population (Figures 3C and S5B).

### Computations that may underlie phase modulation

The neural sensitivity to phase within cycle is surprising, particularly for the random cycle sequences. We describe here a simple model that shows phase modulation for random cycle condition, starting with the responses of a population of frequency-sensitive neurons.

The models consist of modules, each defined by a period P. Each module averages the outputs of multiple chains as described in Figure 4A. The chains of each module have the same P but different sensitivity to frequency.

**Figure 4.**
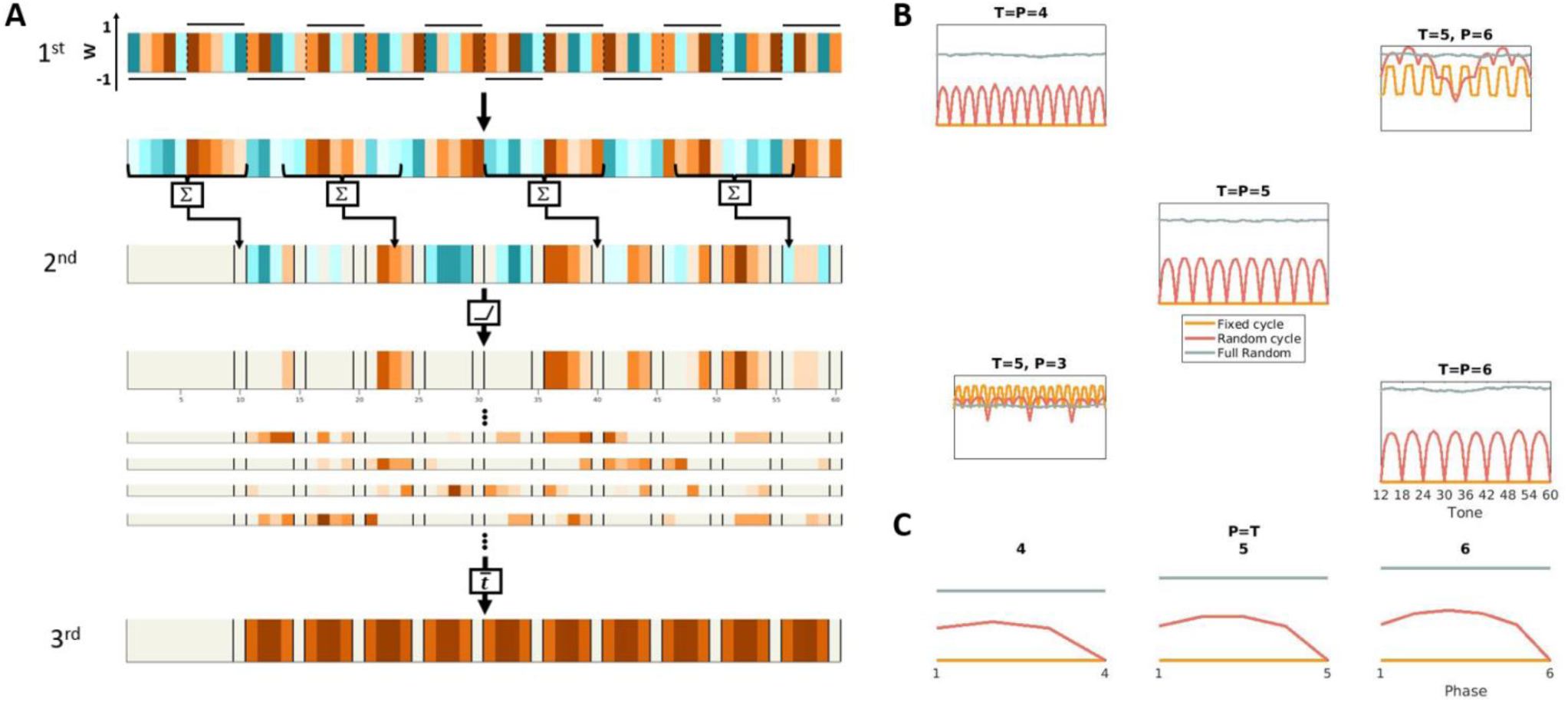
The model A. Schematic explanation of the model. A P=5 chain processes a random cycle sequence with a matching T=5. The color scale represents response strength (dark blue to dark brown, low to high). 1st layer neuron: the top strip represents the frequency- sensitive inputs. Each frequency is associated with a somewhat different response, so that frequency is represented by color. Since the simulations are noise-free, these responses are identical every time a given frequency occurs. The neuron weighs these inputs according to their position in the sequence (the weights are indicated by the black lines around the top strip; the weighted responses are represented in the strip below). The 2^nd^ layer neuron sums of the last 2P outputs of the 1^st^ layer neuron (top strip) and its output is rectified (middle wide strip). The black lines in the third row onwards highlight the positions that are multiples of P. For random cycle sequences illustrated here, when P=T, the output of the 2^nd^ layer neuron is always 0 at these positions. The narrow strips illustrate the output of additional 2^nd^ layer neurons in the same module, which process the outputs of other 1^st^ layer neurons that have different frequency sensitivities. The 3^rd^ layer neuron averages all of the 2^nd^ order neurons, providing the final output of the module. B. Examples of outputs for various P and T values in the fixed cycle (orange), random cycle (red) and full random (gray) conditions. The panels on the diagonal show the outputs in the P=T (4/5/6) cases. The off-diagonal panels show cases with P≠T. To extract the phase functions from the model’s output, we averaged these values over cycles. Importantly, for the P=T cases, the resulting phase functions in the random cycle condition are non-flat, being non-zero at phases 1 to T-1, but having a 0 at the T’th phase.

Figure 4A shows an example of a P=5 chain that processes a random cycle sequence with a matching T=5. The core operation of the module is performed by the 1^st^ and 2^nd^ layer neurons. The 1^st^ layer neurons are driven by the frequency-sensitive inputs (Figure 4A, top line). The frequency-sensitive inputs are weighed by the position of tones along the sequence. The responses to the first P tones are multiplied by -1, then the next P responses by +1, then the next P responses by -1 and so on (Figure 4A, first and second lines). The 2^nd^ layer neuron sums linearly the last 2P outputs of the 1^st^ layer neuron (temporal summation, Figure 4A, third line) and its output is rectified (Figure 4A, fourth line). Finally, the 3^rd^ layer neuron sums the outputs of multiple such chains, with P=5 but differing in the frequency sensitivity of their 1^st^ layer inputs (represented by the thinner rectangles in Figure 4A), providing the output of the module.

The position-dependent weighing of the 1^st^ layer neuron can be replaced by time-invariant filtering using an impulse function whose duration is 2P tones, and which is -1 for the duration of P tones, and then 1 for the duration of the next P tones. While time invariant filtering may be more natural for neural models, some finer features of the model are better apparent when using position-dependent weighing as in Figure 4A, and we prefer to continue describing the results of this version of the model.

Consider such a module with P=T. In the fixed cycle condition, due to the fixed order of the frequencies in all cycles, the positive and negative weighted values cancel each other at every position along the sequence when integrated by the 2^nd^ layer neuron. Therefore, the output of each chain is identically 0, and the module output is identically 0 as well.

In contradistinction, for full random sequences, in general the positive and negative values never cancel each other exactly, and after rectification and averaging over many frequency-sensitive inputs, the output of the module is largely constant as a function of phase and cycle. The model therefore distinguishes easily between fixed cycle and full random sequences.

Importantly, the model also distinguishes random cycle sequences from the fixed cycle and full random sequences. For random cycle sequence, the output of each chain is 0 at times 2T, 3T, 4T and so on because at these times P values with positive sign are summed with the same P values with negative sign (Figure 4A, second and third lines). At other times these sums can be positive or negative depending on the order of the sounds and the frequency sensitivity of the inputs to the chain. Thus, after rectification and summation the output of the module is 0 at times that are multiples of T, and is non-zero at all other times along the sequence (Figure 4A).

Figure 4B shows the average output of the model for varying P and T. For full random sequences, the outputs are non-zero and constant, independent of the nominal T and the P of the module. For P=T (Figure 4B, three larger panels on the diagonal) the output for fixed cycle sequences is identically 0, and for random cycle sequences the output has a periodic sequence of 0s at intervals of T tones. When P≠T (the periodicity of the sequence is different from the characteristic period of the module; Figure 4B, the two off-diagonal smaller panels), the outputs in the fixed cycle and random cycle conditions show fluctuations that reflect the interference between T and P.

To extract the phase functions, we averaged these responses over cycles. Importantly, when P=T, the resulting phase function are non-flat in the random cycle condition, with a 0 at the T’th phase (Figure 4C). The P=T modules show clear preference for randomness, with the responses to the full random conditions largest, the fixed cycle sequences smallest, and the responses to the random cycle sequences intermediate in size. Modules with P different from T may show different sensitivities to randomness. Simulations suggest that when T/P<1.5, the average output of the model is greater for the full random than for the fixed cycle condition, while when T/P>1.5, responses in the fixed cycle condition tend to be higher (Figure 4B).

When estimating phase functions from recorded data, we averaged the responses of the same neuron over sequences composed of many different frequencies, while in the model, the phase functions are derived from averages of many modules, each with its own set of frequency-specific inputs, to the same tone sequence. We argue that varying the sequence (which happens when computing phase functions using experimental data) would achieve the same effect as keeping the sequence fixed but varying the frequency selectivity of the neurons in layer 1 (as we do in the model). Therefore, the phase functions of the model correspond well with the phase functions as estimated from the neuronal responses. In that sense, the model accounts for the existence and strength of phase modulation, particularly in the random cycle condition.

## Discussion

We recorded neural responses to multitone sequences from auditory cortex of awake rats. The tones were presented in three different conditions – fixed cycle, full random, and an intermediate random cycle condition. We show that units in the awake rat auditory cortex are sensitive to the structure of these sequences, and detect even the subtle cycle structure of the random cycle sequences. Furthermore, these units largely align on a continuum of preferences between order and randomness manifested by their different average firing rate in the three conditions.

### Limitations

The study has been run in awake but non-behaving rats. In consequence, there is no control for their state and attention. In addition, the neurons whose responses are documented here are unidentified, and some of the variability between neurons could be related to their type. Finally, the neurons analyzed here are about 10% of the total number of neurons recorded and are therefore spatially sparse. We did not find a clear spatial clustering of the neurons based on their sensitivity to condition, cycle, and phase, so that there is no information available here about the functional anatomy of these effects.

### Differences between responses in anesthetized and awake cortex

As expected from the pattern of results in anesthetized rats, averaged over all units, responses to tones in full random condition were larger than responses to the same tones in fixed cycle condition, with random cycle responses intermediate. However, the average effects hide the large variation between units. While many units showed preference for full random sequences, many other units showed the opposite preference: they responded most strongly to fixed cycle sequences, and least strongly to the full random sequences. Although we didn’t test the same sequences under anesthesia, this result was unexpected: under anesthesia, random two-tone sequences elicited stronger responses than periodic sequences with the same frequency composition in the large majority of neurons^16^.

Similarly, in anesthetized rats, responses tended to decrease along sound sequences^19–21^. In the awake, non-behaving rats, the average population responses did not show significant changes, either increase or decrease, over cycles. In this case too, the average effects masked a large variation between units, and individual units showed responses that tended to increase, remained stable, or had a more complex temporal modulations along the sequence (Figures 2D and S3).

### Rat and human responses to regularity and randomness

Previous human studies have consistently shown higher responses to fixed cycle sequences compared to the full random sequences used in our study^17,18^. Our data provides two clues that may account for the differences between rats and humans.

One clue is based on the finding that in rats there are roughly equal populations of randomness-preferring and regularity-preferring neurons, although there was a small excess of randomness preference expressed in the significant main effects of condition. Since the human signals reflect average activity of large populations of neurons, it could be that in humans there is also a mix of randomness-preferring and regularity-preferring neurons, but the balance is reversed, resulting in an average response that is larger in the regular condition.

Alternatively, the differences may be explained by the time course of the responses to the individual tones. In our data, the population mean firing rates before and after the peak responses to each tone tended to be higher in the fixed cycle conditions compared to the random cycle and full random conditions (Figure 2B). The higher baseline firing rates in the fixed cycle condition may be related to the larger brain signals evoked by the fixed cycle sequences compared to full random sequences in the human studies.

### Sensitivity to sequence structure

Remarkably, some neurons in auditory cortex were sensitive to the position of sounds within cycles (phase sensitivity). About 5% of the sound-responsive units showed significant phase modulation in the fixed cycle and random cycle conditions, a significantly larger fraction than that found in our ‘negative control’ condition, the full random sequences.

The strongest and most consistent phase modulations occurred in the random cycle condition. This was true both at the single unit level and for the averaged population responses. Remarkably, the population phase functions were consistent for all periods (T=4, T=5 and T=6) – roughly monotonically increasing for INC units and monotonically decreasing for DEC units. This result was reproduced in all tested animals and stimulus sets.

The detection of the periodic structure of fixed cycle sequences is not particularly surprising – a periodic sequence of tone frequencies would induce periodicity in the responses. Such periodicity can be detected by autocorrelation or similar computational schemes. Indeed, previous experimental work, even with just repeated segments of white noise, showed strong perceptual effects of stimulus periodicity^22^ for periods in the range tested here (200- 300 ms). Thus, phase sensitivity is particularly remarkable for random cycle sequences, because it shows that the auditory system is capable of detecting the cycles using the subtle cues available in that case. Detecting the cycles of random cycle sequences is a different computational task than in the fixed cycle case, and has not been described before.

### Sensitivity to random cycle sequences can result from a simple computation

We suggest here a simple computational scheme that can distinguish between the different conditions and shows phase modulation for random cycle sequences, mimicking our experimental findings. The model is sensitive to an abstract feature – the potential boundaries between cycles.

The requirement for a counter, triggered by sequence onset, that is used to switch the sign of the weights imposed by the 1^st^ layer neurons every P tone presentations may seem unnatural. As stated in the paper, the counter could be replaced with a time-invariant filter whose impulse response performs the same summation. However, oscillators whose period is consistent with that required here (between 0.4 and 1.2 seconds, about 0.5 to 2 Hz) are ubiquitous in the nervous system, and the entrainment of oscillations at these rates have been implicated in high-level processing of sounds in humans^23,24^ as well as in animals^25,26^. The counter required here could originate from similar mechanisms.

Our model requires the neurons to integrate information over at least two cycles of the sequences, corresponding to 400 ms (8 tones, T=4) up to 600 ms (12 tones, T=6). Previous studies demonstrated that neuronal responses in auditory cortex may depend on a very long history of the stimulation sequence. Thus, it has been shown that responses to oddball sequences composed from two tones increased with increased sequence randomness, requiring sensitivity to at least 40 previous sound presentations^16^. Similarly, a previous study demonstrated that prediction errors in 2-tone sequences are computed relative to predictive models with a long memory (at least 10 sound presentations)^27^. The integration time window required here is longer than that of the single tone response (50 ms), but it is not very long – at 600 ms it is still substantially shorter than the integration windows required to account for the results of previous studies^16,27^.

Previous work also suggested that while memory duration in auditory cortex is long, the representation of the details of the sequence in memory is coarse^27^. The model suggested here also results in a coarse representation of the details of the tone sequence – for example, the temporal summation used here loses substantial amount of information about the order and identity of the sounds.

Why should the auditory cortex care about identifying the random cycle condition relative to the other two? One way to understand this is to consider that the random cycle condition is the only one which has uniquely defined cycles. In the fixed cycle condition, cycles exist, obviously, but there are many choices for them – any successive T sounds would do. In the full random condition, cycles do not exist by definition. Only in the random cycle condition there are uniquely-defined boundaries between cycles. Therefore, it is possible that the higher sensitivity to phase in the random cycle sequences indicates the sensitivity of the auditory cortex to the boundaries of the cycles that compose the sequence.

## Resource availability

### Lead Contact

Further information and requests for resources and reagents should be directed to and will be fulfilled by the lead contact, Israel Nelken.

### Materials availability

This study did not generate new unique reagents.

### Data and code availability

Data and code are available through the lead author

## Acknowledgments

This research was supported by grant 1126/18 of the Israel Science Foundation, grant 2016688 of the NSF-BSF-NIH Computational Neuroscience (CRCNS) program, and ERC Advanced Grant 340063 (project RATLAND). We thank Ana Polterovich for her help in setting up the experimental system, Mousa Karayanni for his help with the experimental system and preprocessing, and the rest of the lab members for their on-going support.

## Author contributions

Conceptualization, methodology, funding acquisition, and supervision: IN. Data acquisition: MMJ. Analysis, methodology, investigation and visualization: SB.

## Methods

### EXPERIMENTAL MODEL AND STUDY PARTICIPANT DETAILS

The experiments were carried out in accordance with the regulations of the ethics committee of The Hebrew University of Jerusalem. The Hebrew University of Jerusalem is an Association for Assessment and Accreditation of Laboratory Animal Care (AAALAC) accredited institution. Four adult female Sprague Dawley rats (Envigo LTD, Israel; weight: at least 200 gr) were used for the experiments described here. All efforts were taken to create a low-stress, rat-friendly living environment enabling experimental animals to freely express their innate behaviors. At their arrival, the rats were housed in the SPF room in which the experimental setup was situated and where all experiments took place. The animals were housed in pairs until implantation and then individually in neighboring cages. The temperature (22 ± 1 °C) and humidity (50 ± 20%) in the room were controlled and the room was maintained on a 12-h light/dark cycle (lights on from 07:00 to 19:00).

### METHOD DETAILS

#### Surgical procedure

The rats were initially anesthetized in an induction chamber with sevoflurane (8% in air, Piramal Critical Care Inc., Bethlehem, PA, USA). Their heads were shaved and they were placed in a stereotaxic instrument with a mask for gas anesthesia (David Kopf Instruments, CA, USA). Sevoflurane concentration was slowly adjusted to 3.7%, and maintained at this level throughout the surgery. The surgical level of anesthesia was verified by a lack of pedal-withdrawal reflex and by the maintenance of a slow and steady breathing rate. Body temperature was controlled with a rectal probe by a closed loop heating system. The eyes were protected with a thick layer of Vaseline and the skin on the head was disinfected with povidone-iodine solution (10%, equivalent to 1% iodine, Rekah Pharm. Ind. Ltd., Holon, Israel). To prevent postoperative pain, rats received subcutaneous injections of Carprofen 50mg/ml (12 mg/kg; Norocarp, Norbrook Laboratories Limited, Newry, Co. Down, Northern Ireland), during and after the surgery, and in the first few days postoperatively if any symptoms of pain were identified.

A 1.5–2 cm longitudinal cut of the skin on the head was made and the dorsal surface of the skull was exposed. The connective tissue covering the bones was removed and the bones were cleaned with sterile saline. Then the surface of the bones was treated with a 15% hydrogen peroxide solution (Sigma Aldrich Inc., St. Louis, MO, United States) and the area was flushed with sterile saline after 10–20 s. When the surface of the skull was clean and dry, a reference point for the entrance of the recording electrodes was marked to target the auditory cortex at the following coordinates: anteroposterior (AP) = -5.1 mm, mediolateral (ML) guided by landmarks on the lateral surface of parietal and temporal bones, and dorsoventral (DV) = 3.2 mm. Six supporting screws were mounted in the skull. One screw soldered to a ground wire was placed in the left frontal bone. The screws were fixed to each other and to the bone with resin and then with acrylic dental cement (Super-bond C&B, Sun Medical, Moriyama, Shiga, Japan; Coral-Fix, Tel Aviv, Israel), forming the base of the implant. The ground wire was connected to a small female connector (853 Interconnect Socket, MILL-MAX MFG. CORP., New York, United States) embedded in the cement in the front of the implant base, creating an easy, low impedance connection to the ground wire during recordings. A thin polyimide tube was placed on the skull vertically above the projected electrode implantation site and cemented together with the rest of the implant base. The wounds were cleaned and treated in situ with antibiotic ointment (synthomycine, chloramphenicol 5%, Rekah Pharm. Ind. Ltd., Holon, Israel) and Bismuth subgallate (Dermatol, Floris, Kiryat Bialik, Israel). The skin was sutured (Nylon, Assut sutures, Corgémont, Switzerland) in the anterior part of the implant with one or two sutures to stretch the skin around the base of the implant. Rats received an intraperitoneal injection of Enrofloxacin antibiotic 50 mg/ml at a dose of 15 mg/kg diluted with saline for a total volume of 1 ml (Baytril, Bayer Animal Health GmbH, Leverkusen, Germany). For the first two days after surgery, Meloxicam (Loxicom 1.5 mg/ml, Norbrook, Newry, Northern Ireland, United Kingdom) was dissolved in palatable wet food and served to the rats in their home cages (0.6 mg in one portion given every 24 h). Rats were allowed at least 2–4 weeks of recovery post-surgery before starting the next procedure. After surgery, the animals were housed individually to prevent damage to the implants.

When the wounds had completely healed we implanted the recording platform. The induction, animal preparation, and care during the surgery were the same as above. The dental cement above the implantation site marked by the polyimide tube was removed using a dental drill until the skull was exposed. The craniotomy was performed by drilling, and the dura was gently resected (0.3–0.5 mm long incision). Electrodes were inserted into the brain tissue using a micromanipulator (to a depth of 1.1–1.5 mm below the brain surface) at a rate of 100 μm/min. Neural activity was monitored during insertion to identify the optimal depth. The craniotomy was sealed with elastic silicone polymer (Duragel, Cambridge Neurotech, United Kingdom) and the electrodes were fixed to the base of the implant with acrylic dental cement. To prevent postoperative pain, the rats received a subcutaneous injection of Carprofen 50 mg/ml (5% W/V) at a dose of about 12 mg/kg. For the first two days after surgery, Meloxicam dissolved in palatable wet food was served to the rats in their home cages (0.6 mg in one portion given every 24 h).

#### Electrophysiology

We used a headstage designed for wireless recordings with Neuropixels probes, together with a logger docking station (LDS_1) for downloading the data from the micro SD card on which they are stored during the experiment (SpikeGadgets, CA, USA). Although the electrodes were fixed throughout the recording sessions, the set of responsive neurons changed substantially across days, with active neurons appearing and disappearing from one day to the next. We therefore consider each recording session as an independent data set.

The recordings were processed through the spike sorting program kilosort3. The results were further analyzed without manual curation, so that all responses recorded using the Neuropixels electrodes are considered as multiunit clusters rather than well-separated units.

#### Multitone sequences

Every sequence consisted of 60 tones, 50 ms long with a linear rise/fall time of 5 ms, presented without gaps between them. The total duration of each sequence was 3 seconds, and they were played every 4 seconds. In all sequence sets, we started by selecting T frequencies (T=4, 5, 6) from a table of 20 possible values (range: 4-40 kHz, 6 tones/octave). From each combination of T sounds we generated sequences for all three conditions. Fixed cycle sequences were generated by selecting a random permutation of the T sounds and then repeating it until the end of the sequence. Random cycle sequences were generated by selecting a new random permutation for every T sounds. Full random sequences were generated by selecting a random permutation of all 60 sounds, consisting of T frequencies each of which repeated 60/T times. In each recording session, the sequences in the block were presented in a fresh random order.

We wanted to study the modulation of the neuronal responses by phase. In a specific sequence, the responses could show an apparent phase dependence as a consequence of the specific order of the frequencies due to neuronal frequency tuning. This was especially so in the fixed cycle condition because in this condition the same frequencies are presented in the same phases at all cycles. We controlled for frequency sensitivity in three ways. We used sequences with different choices of the T frequencies; for every choice of T frequencies, we generated sequences of all types (Figure 1A); and for every sequence used in the experiment, we added T-1 sequences for control and balance in each condition (Figure 1B). In the fixed cycle condition, for every choice of T frequencies we generated one sequence and then added T – 1 cyclic permutation of that sequence, such that each one of the T selected tones appeared at all the possible phases while its position relative to the other tones remained the same. Thus, while the periodic structure of the stimuli in reg sequence was expected to induce a periodic component in the neuronal responses, this periodicity shifted circularly and therefore was expected to result in a constant average firing rate as a function of phase. In the random cycle condition, for every sequence we added T – 1 sequences that contained the same frequencies but were independently randomly permuted for every cycle. In the full random condition, we added T – 1 random permutations of the entire sequence.

##### T = 4/6

We generated these sequences using 20 random combinations of T frequencies out of the table of 20 frequencies, with the requirement that each frequency appeared in T of these 20 combinations. Each of these 20 combinations of T frequencies was used to generate T sequences for every condition as described above. Each sequence was played twice within a session. Thus, the number of sequence presentations in each condition was 20·T·2 (160 for T=4 and 240 for T=6).

T=4 and T=6 sets were always played in the same recording sessions, so that the neural data for these two sets were recorded from the same units. The block of T=4 sequences were presented before the block of T=6 sequences.

##### T = 5

For T=5 we used several sets of sequences.

##### The main set

In this set we generated sequences using 16 random combinations of 5 frequencies out of the table of 20 frequencies, with the requirement that each frequency appeared in 4 of these 16 combinations. Each of these 16 combinations of 5 frequencies was used to generate T sequences for every condition as described above. Each sequence was played once within a session. Thus, the number of sequences played for each of the conditions was 5·16=80. The data for T=5 in the Results section are all from recording sessions using this set.

##### Set1 and Set2

In set 1 we generated sequences using only 2 combinations of 5 frequencies. The frequencies were selected to be equidistant in each combination, and covering roughly the same, large, frequency range ([4, 6.5, 10.55, 17.13, 27.81], [5.1, 8.3, 13.44, 21.82, 35.43] kHz). Each of the two combinations was used 10 times to generate 5 sequences of each condition, as described above. Each sequence was played once within a session, for a total of 2·10·5=100 sequences in each condition. Thus, this is the set in which the frequencies were the most balanced, but with little variability in the frequency composition of the different sequences.

Set 2 used the same two frequency combinations of set 1 with one additional combination ([4.52, 7.33, 11.91, 19.33, 31.39] kHz). Otherwise, the construction of the sequences was the same as for set 1. Each sequence was played once within a session, for a total of 3·10·5=150 sequences in each condition.

##### Set3

We selected 10 random combinations of 5 frequencies, without any attempt at balance as in the other sets. From each of these combinations we generated 5 sequences for each condition as described above. Each sequence was played once within a session. The number of sequences played in the block for each of the conditions was 5·10=50.

Data from recording sessions using Set1, Set2 and Set3 are shown in the supplementary data figures.

### QUANTIFICATION AND STATISTICAL ANALYSIS

Statistical analysis was performed using MATLAB (The MathWorks, Inc., Natick, MA, USA).

#### Exclusion criteria for trials

Animal movements caused sometimes large artifacts on many channels simultaneously.

Recordings with artifacts usually showed positive and negative voltage peaks with values that exceeded the physiological range. Trials that had artifacts of this type were excluded from the analysis. In order to identify them we summed the voltage at each moment in time over all channels. For each trial we computed the variance over time of the summed voltage trace (termed ‘x’ below) and the maximum absolute value (termed ‘y’ below). We rejected trials that had log(y) > 18.3 -0.322log(x). This criterion resulted in the rejection of about 2% of the trials (81,235/83,210 trials were included in the analysis).

#### Auditory units

In order to identify auditory units, we used an inhomogeneous Poisson likelihood ratio test. We used the firing rate during the 50 ms preceding the stimulus and compared it to the firing rates in 5 bins of 10 ms covering the first 50 ms of the sequence (first tone presentation). For each 10 ms bin, we computed the Poisson tail probability of observing that spike count assuming the expected spike count from the pre-stimulus period, and units were considered auditory units when at least one of the bins had p<0.01. We performed one-sided tests for the upper tail of the distribution (“INC” units) and for the lower tail of the distribution (“DEC” units). To avoid false detections when units had a very low firing rate, one count was added to the expected spike count preceding the stimulus. A total of 12,363/58,602 units were found to be auditory based on this criterion.

#### Exclusion criteria for units

Units that fired in less than 10% of the trials were not included in the analysis. This criterion resulted in exclusion of about 26% of the units that were identified as auditory units (9,165/12,363 units remain after this exclusion).

Additionally, we excluded unstable units. We used two stability measures. (1) For each unit, we calculated the variance over time of each trial and averaged these over trials. This was divided by the average of the variance over trials at each point in time (1 ms bins). (2) For each unit, we cross-correlated its spike counts per trial with the spike counts per trial of all other units. We then computed the ratio of the mean to the standard deviation of these values. Stable units were selected to be with y+121x > 122 (where x is the value for (1) and y is the value for (2); This line was selected based on manual checks of the stability of selected units). Measure x was small when the firing rates changed substantially across trials (the denominator was large in these cases). Measure y was small when the response profile of the unit was very different from that of all other units. This happened usually when units appeared or disappeared during a block. The test tended to make both x and y large.

Next, we further examined the stability of the recordings. We computed the spike counts in each trial. We correlated these values with those of units previously determined as unstable, and for each unit we extracted the maximum absolute value of these correlations (termed ‘x’ below). In addition, for each unit, we sorted the spike counts (ignoring those emitted during the first and the last 20 trials, to avoid large differences due to spike rate adaptation). The sorted trials were smoothed (window size: 30 trials). Finally, we calculated the average values of the 20 trials with the highest firing rate (mean_max), the 20 trials with the lowest firing rate (mean_min), and computed (mean_max – mean_min)/mean_max (termed ‘y’ below). The final selection of stable units required x+2.4y < 2.3. As before, the separation line was determined by checking visually the stability of a selected set of units. Here, x represented correlations of the sequence of spike counts with those of units that were already removed from the dataset; y was small when the total spike counts across trials didn’t change much. The criterion tended to make both values small.

These two stability criteria resulted in the rejection of about 26% of the units that remained after the first exclusion (6,756/9,165 units from all sets were included in the analysis, 4404 units from the main data set).

|  | Main Data |  | Additional Data<br>T=5 |  |  |
| --- | --- | --- | --- | --- | --- |
|  | T = 4/6 | T=5 | Set1 | Set2 | Set3 |
| **All units** | 24698 | 16309 | 8281 | 6945 | 2369 |
| **Auditory units included in the analysis** | 2426 | 1978 | 962 | 939 | 451 |
| **INC units** | 1200 | 1350 | 479 | 587 | 382 |
| **DEC units** | 1226 | 628 | 483 | 352 | 69 |
| **Units from each rat** | Rat 1: 1289<br>Rat 2: 1137 | Rat 1: 1231<br>Rat 2: 747 | Rat 1: 962 | Rat 2: 939 | Rat 3: 397<br>Rat 4: 54 |
| **Recording sessions** | Rat 1: 35<br>Rat 2: 7 | Rat 1: 14<br>Rat 2: 5 | Rat 1: 14 | Rat 2: 7 | Rat 3: 5<br>Rat 4: 2 |

#### Statistical analysis

To study the dependence of neural responses on condition, cycle and phase, linear mixed effects models (MATLAB function ‘fitlme’) were fitted to the number of spikes emitted in 50 ms time windows, corresponding to the durations of the individual tones in the sequences. The fixed factors were the population means, while the unit-specific contributions were estimated by using random factors consisted of the unit-specific deviations from the population mean (random intercepts as well as random slopes).

#### Testing the effect of condition

In order to study the dependence of neural responses on the condition (fixed cycle, random cycle and full random), we modeled the responses to each tone as a sum of contributions of condition and frequency sensitivity factors.

Frequency sensitivity was modeled using two basis functions, each of which consisted of a profile of sensitivity over the 20 frequencies composing the common frequency table we used. The generation of these basis functions is described below.

The Wilkinson notation for the model is:

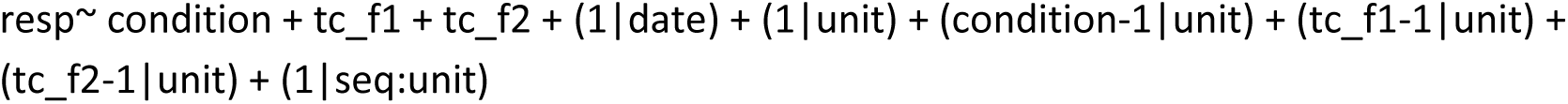

Here, condition is the type of the sequence (fixed cycle, random cycle or full random), tc_f1 and tc_f2 are the two basis functions for frequency profiles. These main effects reflected the population mean. Random slopes included unit-specific contributions for each condition and unit-specific contributions to the weights of the frequency profiles. Random intercepts included a session-specific intercept (date), unit-specific intercepts that accounted for the mean response of the individual units to all tones, and intercepts that accounted for the mean response of each unit to all sequences with the same tone composition (seq:unit).

#### Testing differences between conditions in short time windows

For each unit, we summed the spikes during the first 10 ms and the last 10 ms of the sound for the baseline responses, and the spikes during the 30 ms between them for the peak responses. For testing the effects of condition in each of the time windows, we first computed the mean responses of each unit for each condition in that time window, and then subtracted their mean.

We were interested in the difference between the different conditions, and therefore tested the observed data using a permutation test. In order to get the null hypothesis distribution we permuted the average responses in the three conditions within each unit. We then compared the distribution of the response differences for each pair of conditions with the observed data (Table 4).

#### Testing the modulation by phase and by cycle at the population level

To study the dependence of average responses on cycle and phase we fitted LME models for each condition separately. We decomposed the responses to each tone into a sum of contributions of frequency sensitivity, cycle and phase.

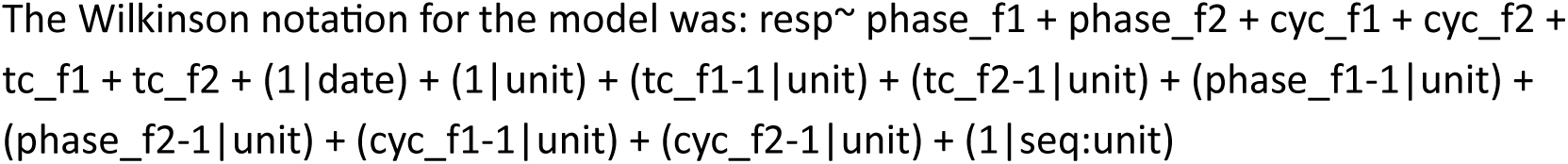

Here, phase_f1 and phase_f2 are two basis functions for phase profiles, cyc_f1 and cyc_f2 are two basis functions for cycle profiles. Their derivation is described below. The other terms were defined above.

#### Testing phase modulation at the unit level

To study the dependence of neural responses on phase within cycle we fitted LME models for each unit and each condition separately. We decomposed the responses to each tone into a sum of contributions of frequency sensitivity, cycle and phase.

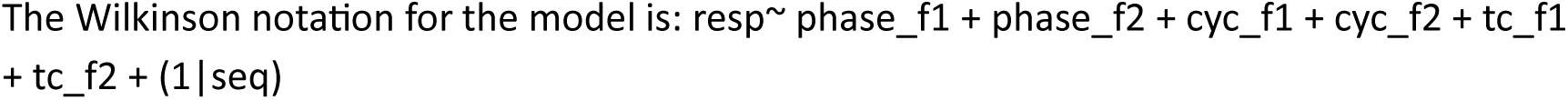

Here phase_f1 and phase_f2 are the two basis functions for phase profiles, cyc_f1 and cyc_f2 are the two basis functions for cycle profiles, tc_f1 and tc_f2 are the two basis functions for frequency profiles, and as above seq identifies sequences with the same frequency composition.

To test the significance of the phase modulation of each unit in each condition, we used permutation tests. First, we computed the likelihood ratio between a model that included the two basis functions for phase, and a model that didn’t include them. We then estimated the distribution of the LR under the null hypothesis of no effect of phase using surrogate data. To generate the surrogate data, for each cycle in each trial, we jointly permuted the values of the two basis functions. This procedure effectively permuted the phase of the responses, while keeping the frequencies and the cycle identity as in the original data. Thus, in the surrogate data, the true phase modulation was flat by design. We then recomputed the model for the surrogate data, with and without the phase basis functions, obtaining the likelihood ratio for the two. This process was repeated 1000 times, resulting in a distribution of LRs for the surrogate data. Finally, we compared the true-data LR with the distribution of the LRs from the surrogate data. We considered phase effects to be significant when p<0.01 in this permutation test.

#### Basis functions

To improve convergence and stability of the statistical models, we described the effects of frequency, cycle or phase as the sum of two basis functions. Each of these basis functions was a profile over the corresponding parameter. The population/unit dependence on that parameter was modeled as a weighted sum of these two basis functions.

As an illustration, Figure S4A shows the first basis functions for cycle for all conditions and data sets. For T=5, for example, these basis functions are profiles over cycles 2-12; a unit would have a high positive weight on this basis function when its response decreased gradually along cycles and a high negative weight when its responses increased gradually along cycles.

The basis functions consisted of the first two principal components determined from the responses a subset of strongly-responding neurons. These were 10% of the units in each recording session that were included in the final data and had the highest firing rate. For example, to determine the frequency basis functions, the responses of each of the selected units were averaged separately for each tone frequency, and the first two principal components were derived from this set of frequency profiles (using the MATLAB function ‘svd’). The cycle (phase) basis functions were derived similarly, based on the average responses to each cycle (phase) of the same units.

---

**Figure S1.**
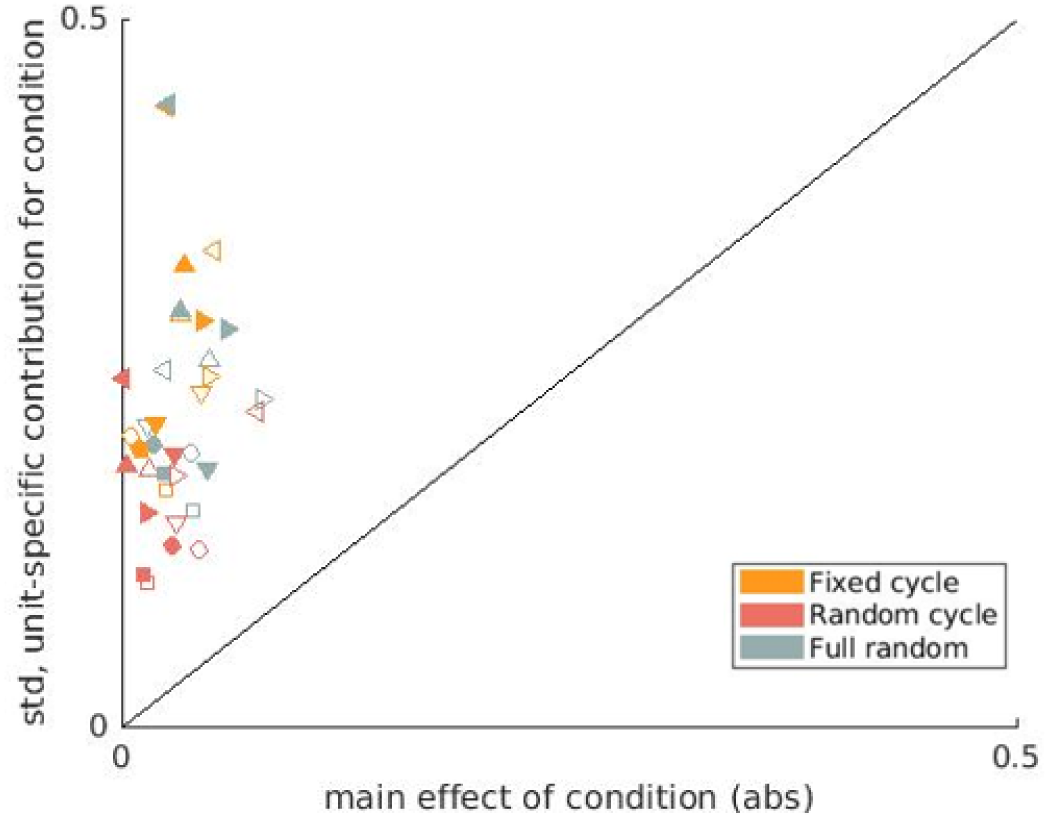
The main effect of condition is substantially smaller than the variability between units The standard deviations of the unit-specific contributions (ordinate) are plotted against the corresponding absolute value of the main effect of the same condition in the model (abscissa). Both are parameters of the same LME model. The main effects correspond to the population average of the effect of condition, while the standard deviation of the unit-specific contributions reports its variability across units. Each point corresponds to one condition from one dataset, and each dataset provides 3 points to the scatter. The figure shows all data sets. ● - T=4, ▲ - T=5, ▀ - T=6, ▼ - Set1, ► - Set2, ◄ - Set3, filled and empty shapes are for tests of the INC and DEC units respectively. See also Figures 2B,C.

**Figure S2.**
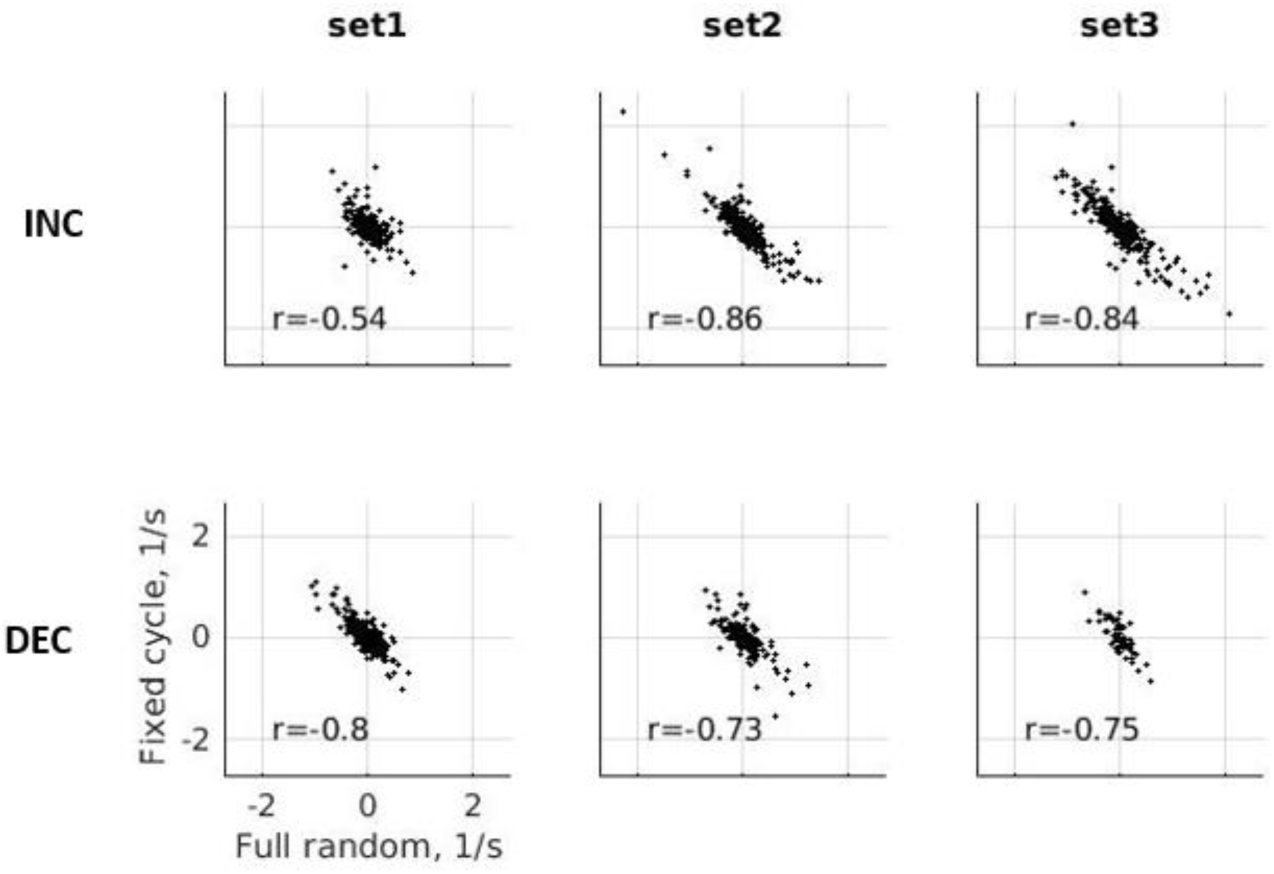
Unit-specific contributions for fixed cycle and full random conditions Scatter plots of the unit-specific contributions in the fixed cycle and full random conditions for Set1, Set2 and Set3 (see Methods for details). The correlation coefficients are presented at the bottom of each panel. See also Figure 2C.

**Figure S3.**
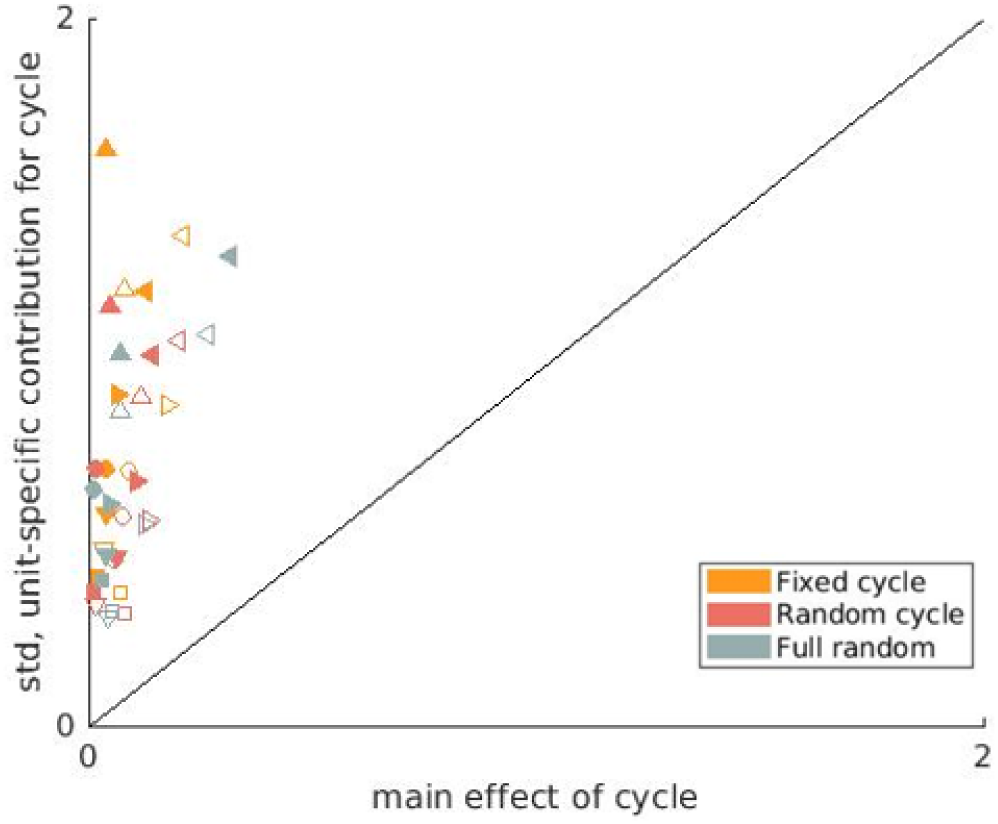
The main effect of cycle is much smaller than the variability between units Same conventions as in Figure S1. See also Figure 2D.

**Figure S4.**
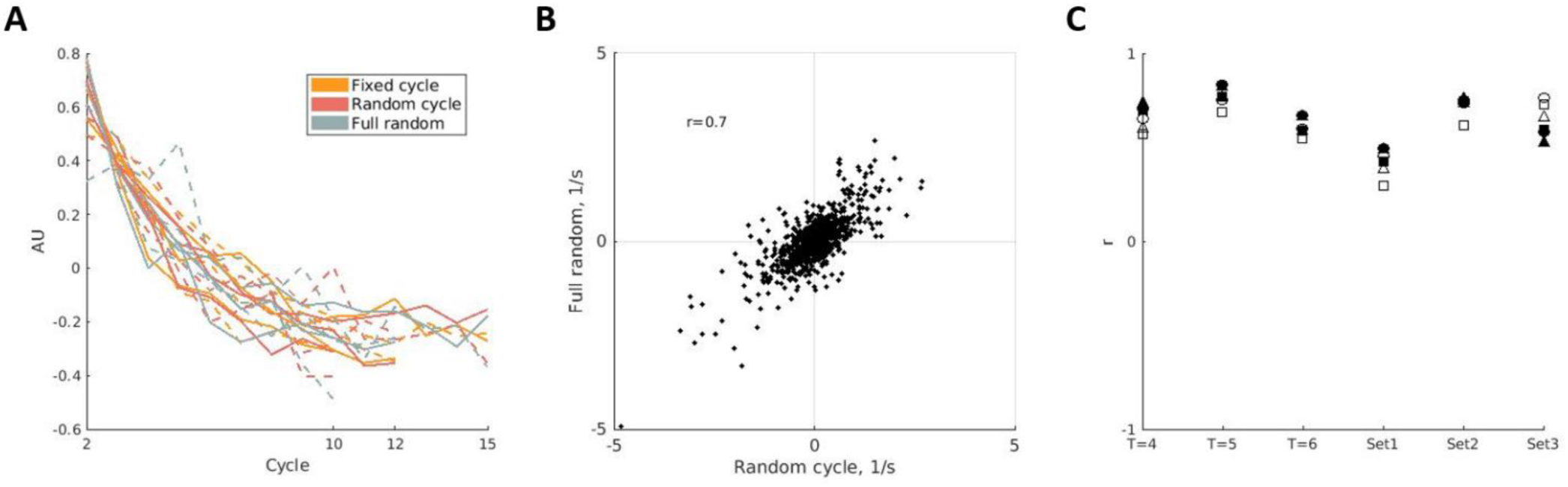
Effects of cycle are similar in all conditions A. The effects of cycle were estimated using a combination of two basis functions (see Methods for details). The first basis functions for cycle were similar across sets and conditions. Continuous line - INC units, dashed line - DEC units. All datasets are represented here. B. Example of scatter plot of unit-specific contributions for the first basis functions of cycle for random cycle sequences (abscissa) and full random sequences (ordinate) of T=4 INC units. C. Correlations between the unit-specific contributions for the first basis functions of cycle in the three conditions. ● - correlations between fixed cycle and random cycle, ▀ - fixed cycle and full random, ▲ - random cycle and full random; Filled shapes- INC units, empty shapes- DEC units. See also Figure 2D.

**Figure S5.**
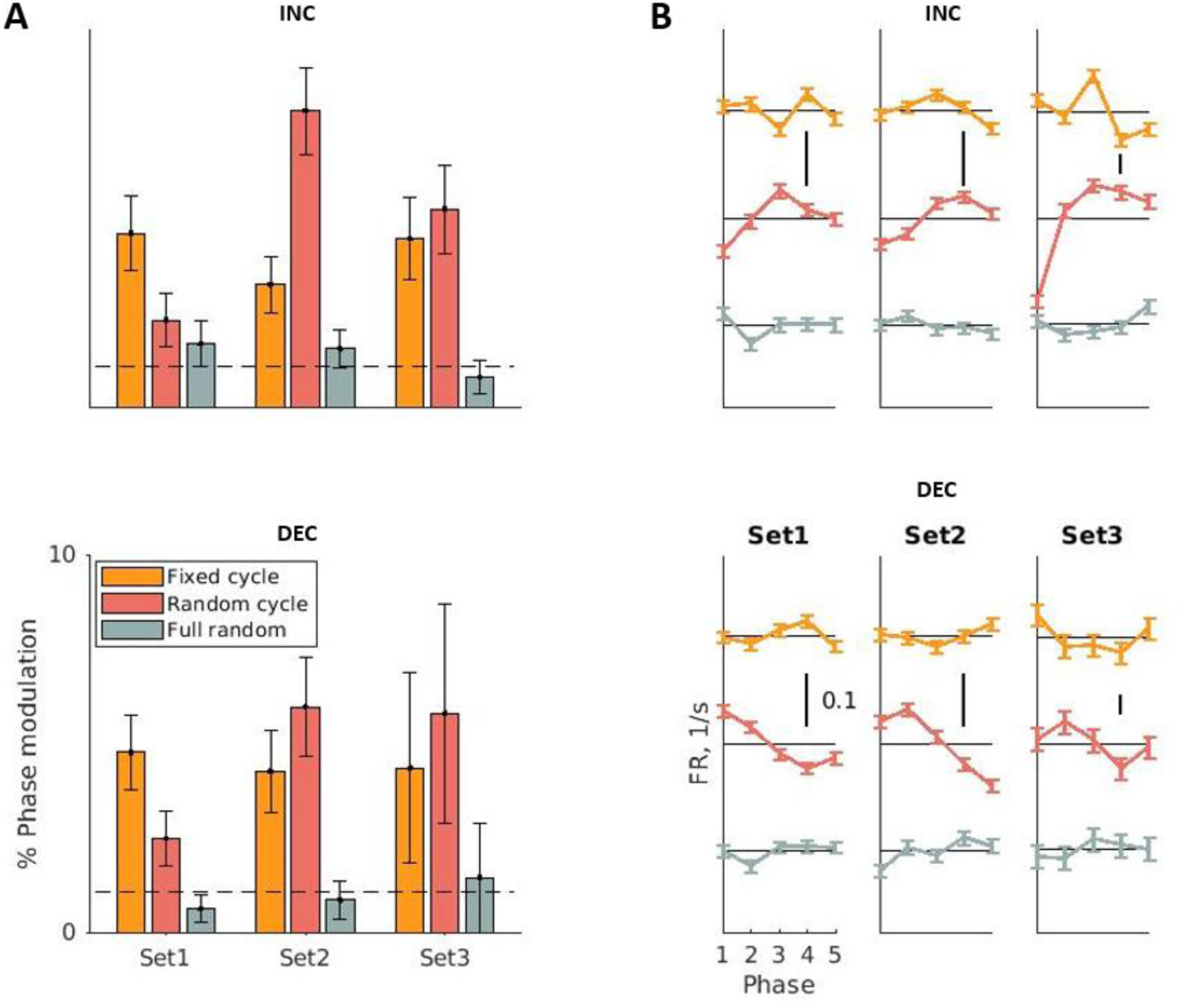
Dependence on the phase in the cycle E. The percentage of units that showed significant phase modulation under each condition for Set1, Set2 and Set3, separately for INC and DEC units. The dashed line marks the expected value under the null hypothesis. See also Figure 3B. and Figure S6. F. The mean population phase functions for the three datasets. See also Figure 3C and Figure S7.

**Figure S6.**
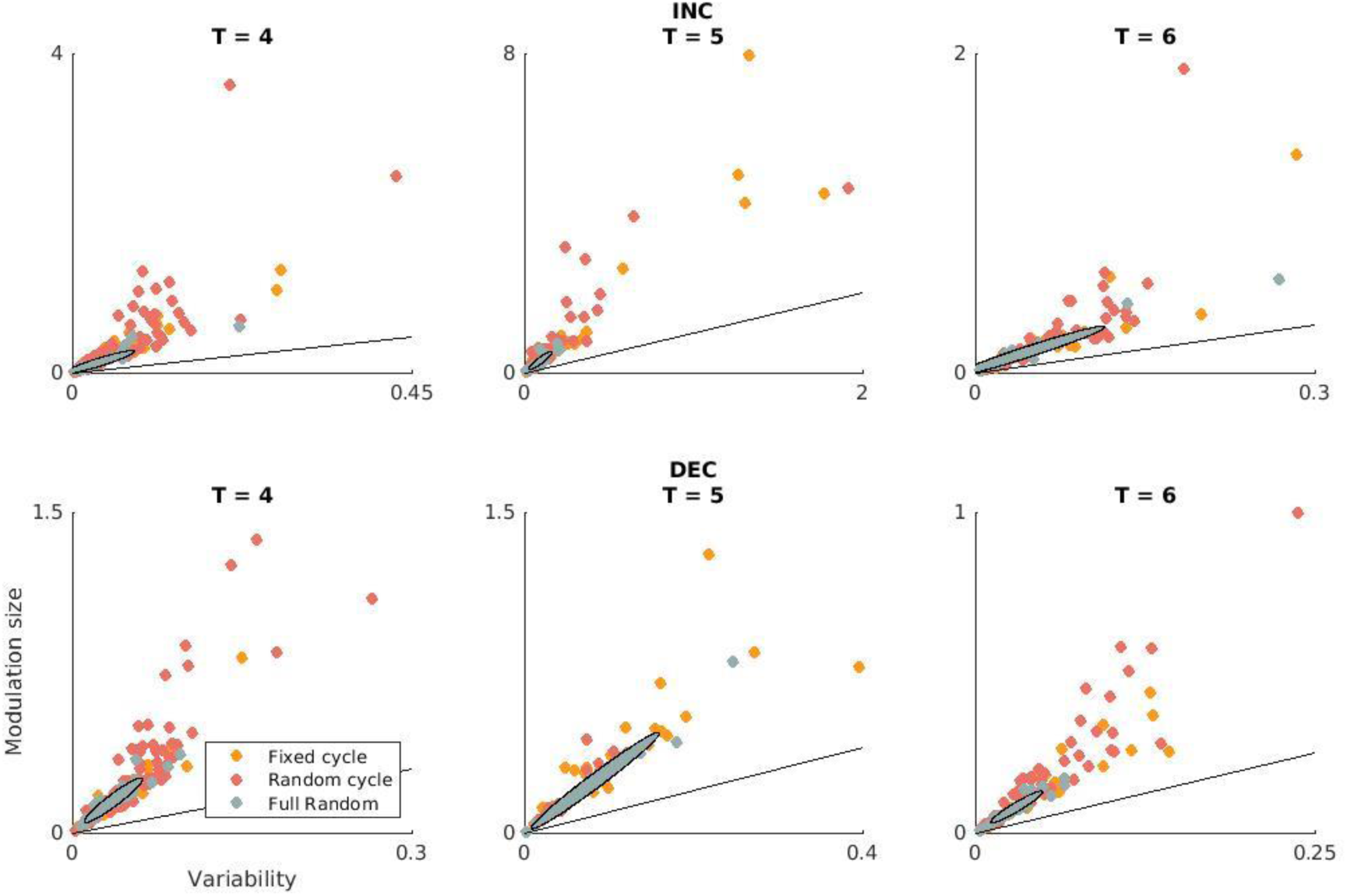
Effect sizes of significant phase functions We calculated for each unit the standard deviation of its phase function (modulation size). Next, we calculated the variance of the responses in each phase, averaged these variances, and then divided by the number of responses used to compute each variance and took the square root (average standard error of the mean, variability). Only units with significant phase modulation are used in these plots. Each dot corresponds to one unit in one condition. The ellipses were derived from the full random condition cases in these plots; they are derived from their covariance matrix and correspond to 1 standard deviation of the plotted values. While the ellipses indeed cover most of the significant cases in the full random condition, there are a large number of cases in the fixed cycle and random cycle conditions outside them. See also Figure 3B.

**Figure S7.**
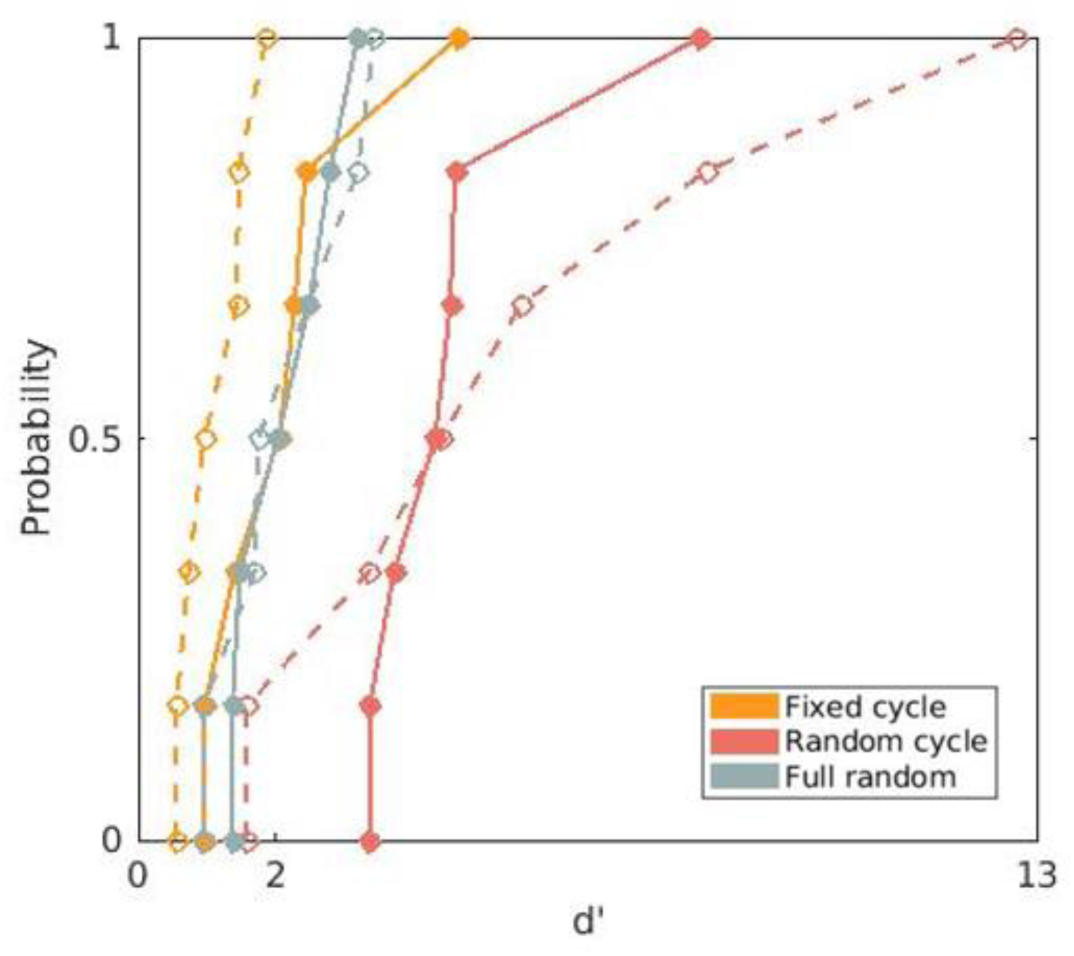
Mean phase effect size To estimate the effect size of the mean phase modulation averaged across all units, cycles, and frequency choices, we calculated for each set the standard deviation of the population mean phase functions (signal). Next, we calculated the average standard error around the phase function, by computing the variance of the unit-specific phase functions in each phase, dividing by the number of units, then averaging over phases and taking the square root (noise). The effect size is the ratio of the signal to noise. The figure shows the cumulative distribution functions of the effect sizes computed in all sets. Continuous line - INC units, dashed line - DEC units. See also Figure 3C and Figure S5B.

